# Visual Continuous Recognition Reveals Widespread Cortical Contributions to Scene Memory

**DOI:** 10.1101/234609

**Authors:** Timothy M. Ellmore, Chelsea P. Reichert, Kenneth Ng, Ning Mei

## Abstract

Humans have a remarkably high capacity and long duration memory for complex scenes. Previous research documents the neural substrates that allow for efficient categorization of scenes from other complex stimuli like objects and faces, but the spatiotemporal neural dynamics underlying scene memory are less well understood. In the present study, we used high density EEG during a visual continuous recognition task in which new, old, and scrambled scenes consisting of color outdoor photographs were presented at an average rate 0.26 Hz. Old scenes were single repeated presentations occurring within either a short-term (≤ 20 seconds) or longer-term intervals of between 30 sec and 3 minutes or 4 and 10 minutes. Overall recognition was far above chance, with better performance at short- than longer-term intervals. A group ANOVA found parietal and frontal ERPs discriminated the three scene types as early as 59 ms after stimulus onset. Parietal ERPs were greater for old compared to new scenes by 189 ms, while fronto-temporal ERPs were greater for new compared to old scenes by 194 ms. For old scenes presented within longer-term intervals, parieto-temporal and centro-frontal ERPs were greater by 228 and 355 ms respectively compared to old scenes presented within a short-term interval. Supervised machine learning exhibited above-chance decoding of scene type by 275 ms. Single-subject BOLD-fMRI showed greater activity for old scenes across frontal, parietal, and temporal cortex. These converging findings show that a widespread network including parietal, frontal, and temporal regions supports short- and long-term scene memory.

**Significance Statement:** The ability to recognize a scene as novel or familiar is critical for basic cognition. Scene recognition plays an important role in episodic memory because it helps us quickly establish place, a first step in recalling where previous events occurred. Short-term recognition supports our ability to detect changes in the immediate environment, an ability critical to survival. Scene recognition after a longer-term interval is often the essential cue for retrieving autobiographical memories. Previous behavioral studies demonstrate high capacity and long duration scene memory. Neural studies have identified the brain regions that support scene-specific processing. The present study extends this research by filling a gap in understanding how distributed spatiotemporal patterns of neural activity support short- and long-term scene memory.

## Introduction

Humans have a remarkable capacity for remembering complex visual information. Early behavioral studies demonstrated that adults and children can recognize large sets of visual stimuli after a single exposure (Shepard, 1967; Standing et al., 1970; Brown and Scott, 1971). While speed of recognition for pictures tends to be slower than for verbal material, reaction times for a range of learning set sizes indicate fast memory search (Standing, 1973). Picture recognition is also highly flexible, with subjects able to discriminate in forced choice paradigms between targets and distractors using perceptual and ecphoric similarity (Tulving, 1981). Early studies of visual memory capacity often mixed objects with travel slides containing complex naturalistic visual scenes. Subsequent research compared encoding of complex scenes with edited versions of scenes that contained a common feature (e.g., a door) and found memory performance for non-edited original photographs was close to 85%. When scene details were removed performance dropped by as much as 20%, suggesting that visual details in scenes contribute positively to long-term memory (Vogt and Magnussen, 2007).

Other findings support that subjects successfully maintain detailed representation of thousands of images (Brady et al., 2008). When the number of exemplars from different categories is controlled for during the study of large picture sets, the capacity to remember visual information in long-term memory varies more with conceptual structure than perceptual distinctiveness. Images from object categories with conceptually distinctive exemplars show less interference as the number of exemplars is increased (Konkle et al., 2010). High capacity picture memory would appear to be at odds with the traditional view that working memory capacity is limited to three or four items. The ability to recognize complex images after short retention intervals would seem to require a larger capacity temporary store, especially if complex details are used. When maintenance using a rehearsal strategy is prevented by using rapid serial visual presentation, memory capacities of up to 30 retained pictures for 100 item lists are obtained, which suggests humans have a larger capacity temporary memory store when proactive interference is minimized (Endress and Potter, 2014).

Scalp EEG has been used to demonstrate fast, parallel processing of complex scenes. In a go/no-go task in which subjects must determine whether a briefly presented scene contains an animal or not, a frontal event related negativity develops on no-go trials approximately 150 ms after stimulus onset (Thorpe et al., 1996). Event related potentials (ERPs) reflect the visual category of a scene as early as 75-80 ms post-stimulus, but are not correlated with behavior until around 150 ms (Vanrullen and Thorpe, 2001). Subjects are as fast at responding to two simultaneously presented scenes as to a single one (Rousselet et al., 2002) demonstrating parallel processing, but behavior and ERPs suffer a processing cost when up to four scenes are presented simultaneously (Rousselet et al., 2004). For biologically relevant scenes, fronto-central ERPs begin to diverge from other stimulus categories around 185 ms after stimulus onset, with a later divergence in parietal regions (Anokhin et al., 2006). Scene recognition is state-dependent and can be modulated by alcohol intoxication (De Cesarei et al., 2006), which reduces early differential ERP activity occurring 150-220 ms when discriminating targets from non-target distractors. An early marker of scene-specific processing was found in a recent study which reported that the first ERP component to evoke a stronger response to real-world scenes compared to other categories is the P2, peaking approximately 220 ms after stimulus onset (Harel et al., 2016).

Intracranial EEG and fMRI studies identify spatiotemporal aspects of scene processing. An intra-cerebral study found early posterior parahippocampal gyrus gamma (50-150 Hz) activity between 200-500 ms when subjects passively viewed scenes (Bastin et al., 2013). Functional MRI activity in both lateral occipital area (LOC) and parahippocampal place area (PPA) can be harnessed to classify scenes accurately. PPA activity confuses scenes that have similar spatial boundaries, while LOC activity confuses scenes that have similar content (Park et al., 2011). Recent work extends the role for occipital place area by demonstrating it can predict pathways for movement in novel scenes (Bonner and Epstein, 2017). It has also been demonstrated recently that humans do not segment a scene into objects but instead use global, ecological properties like navigability and mean depth (Greene and Oliva, 2009). Neural evidence also shows that contrast energy and spatial coherence modulate single-trial ERP amplitudes early (100-150 ms), with spatial coherence influencing later activity up to 250 ms (Groen et al., 2013).

While behavioral studies demonstrate that scene memory is high capacity and long-lasting and neural studies have characterized scene-specific neural processing, the spatiotemporal neural patterns that support scene memory remain to be fully characterized. In the present study, we asked: When do neural patterns distinguish scenes from scrambled perceptual input? How do neural patterns differentiate new from previously presented scenes? And how do neural patterns differ for short- and long-term scene memory?

## Materials and Methods

### Subjects

A total of 29 subjects (mean age 21.21, std. age 2.88, range 18-29, 9 males, 1 left-handed) were recruited between September 2016 and August 2017 by flyers posted throughout the [Author University]. Included in the study were healthy adults between the ages of 18 and 29 with normal or corrected-to-normal vision and the ability to make button presses. Participants were excluded if they did not speak English. Each participant provided written informed consent and completed study procedures according to a protocol approved by the Institutional Review Board of the [Author University]. Participants were compensated $15 per hour for participation. All 29 participants completed the scene memory task during high density scalp electroencephalography (HD-EEG). EEG data from two subjects were excluded because of excessive noise leaving 27 subjects (mean age 21.33, std. age 2.92, range 18-29, 9 males, 1 left-handed). Two additional subjects underwent fMRI (both female, ages 26 and 30).

### Experimental Design

Subjects who underwent HD-EEG completed four 20-minute runs of a visual continuous recognition task (VCRT) during a single session. The VCRT stimuli consisted of color scenes and phase-scrambled scenes (Fig 1a). Scenes were 24-bit color images randomly sampled from the SUN database (Xiao et al., 2010). Only a small fraction (618) of all the SUN database pictures were used in the present study. Care was taken to sample pictures of outdoor scenes with no clearly visible faces. The task was programmed in Visual C++ with graphic presentation optimized by pre-loading as texture maps all stimuli into video RAM using OpenGL. Each stimulus was presented for 1400 ms with jittered interstimulus intervals (ISI). A total of 1228 stimuli were shown during the 80-minute EEG testing session (∼15.35 stimuli per minute). Stimuli consisted of 305 scrambled scenes, 618 new scenes, with 309 of the new scenes subdivided among three old conditions and subsequently repeated one more time 1) within 20 sec, 2) within 30 sec and 3 min, and 3) between 4 min and 10 min. (Fig 1b).

**Figure 1.**
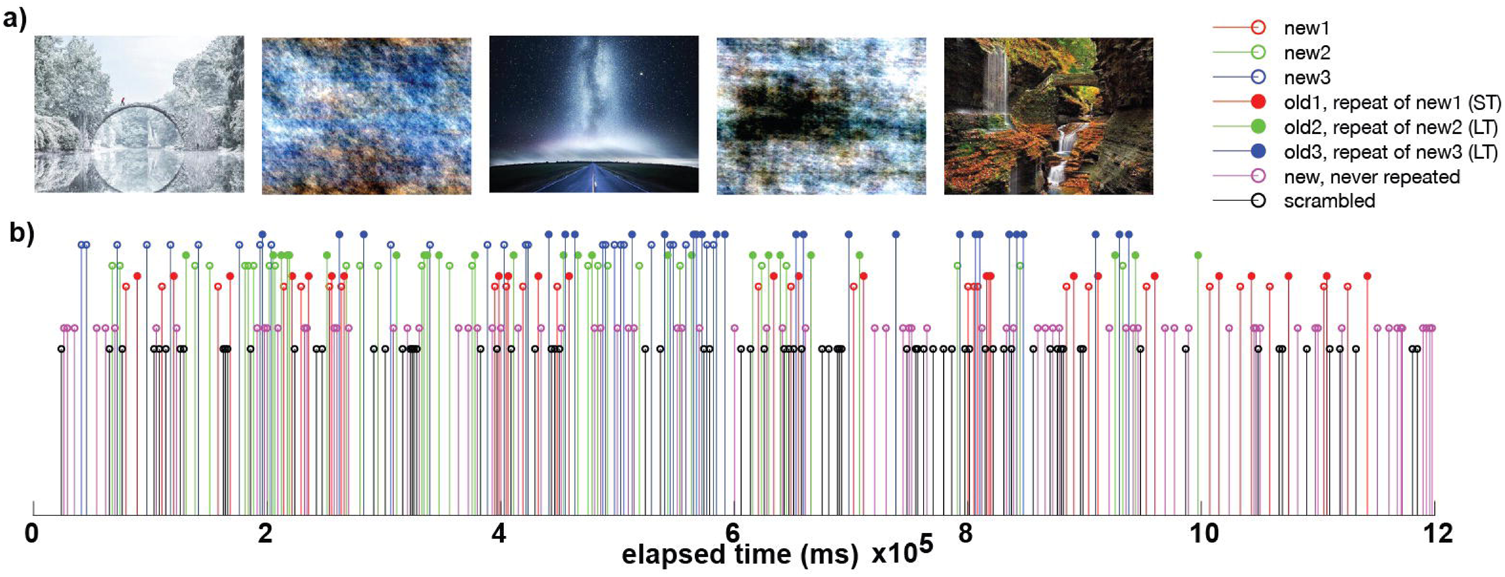
Example Stimuli and Experimental Design. Three example scenes are shown alternating with two example phase-scrambled scenes (a). A stem plot shows 20 minutes of the visual continuous recognition task (b) with each stem representing a single stimulus presentation. The height of the stems is varied simply to aide visualization of the different conditions. Three sets of new scenes to be repeated with old_1_ occurring in the range of shortterm (ST) memory (within 20 sec) and old_2_ and old_3_ occurring in the long-term range (>30 secs after initial presentation). Also displayed are stems representing phase-scrambled scenes and new scenes only presented once during the 20-minute block of VCRT trials.

Each scene was displayed on a 27-inch LED monitor with a refresh rate of 60 hertz (Hz) and a screen resolution of 1920-by-1080. Participants sat 83.5 cm from the monitor and maintained stable viewing using a combined forehead/chin rest. Each scene measured 800-by-600 pixels on the screen, and from the subject’s point of view occupied a horizontal viewing angle of 17.2 degrees and a vertical viewing angle of 12.7 degrees. The EEG recordings took place within a sound-attenuated booth (IAC acoustics) to minimize auditory and visual distractions. Subjects made one of two button (green=old; red=new) responses with their thumb using a fiber optic response device (fORP 904, Current Designs, Inc.) held in their right hand.

### Behavioral Analysis

Analyses of VCRT behavioral data included computing subject accuracy in the form of percent correct in distinguishing old and new scenes. Signal detection analyses were also performed to assess each subject’s recognition ability. A hit was counted when an old scene was correctly classified as an old scene. A false alarm was counted when a new scene was incorrectly classified as an old scene. For each subject, total hits and false alarms were expressed as proportions and used to compute a measure of sensitivity as the difference in standardized normal deviates of hits minus false alarms: d-prime (d’) = *Z(hit rate)* – *Z(false alarm rate)*. The d-prime sensitivity measure represents the separation between the means of the signal and noise distributions, compared against the standard deviation of the signal or noise distributions (Stanislaw and Todorov, 1999).

Overall percent correct and d-prime were based on the ability to recognize scenes as old or new across the four 20-minute blocks. Separate accuracy and d-prime measures were computed for each condition of old: old_1_ (repeated presentation within <20 sec), old_2_ (repeated presentation between 30 sec and 3 min), and old_3_ (repeated presentation between 4 min and 10 min). Average subject response times were also computed for new, old, hits, misses, false alarms, and correct rejections. Repeated measures ANOVAs of accuracy, d-prime, and response times were performed using JASP 0.8.3.1 (https://jasp-stats.org/) with post-hoc tests and Bonferroni multiple comparisons corrections.

### EEG Acquisition

EEG data were sampled at 1 kHz using Pycorder software from 62 scalp locations using an active electrode system with an actiCHamp amplifier (Brain Products). Electrodes were placed at standard locations specified by an extended 10-20 system. The recording ground (Fpz) was located at the frontal midline and the recording reference was located at the left mastoid (TP9) leaving 61 scalp recordings. Two additional channels were designated for left (LOC) and right (ROC) vertical electrooculography (VEOG) recordings for subsequent isolation of eye blink artifacts.

Recordings to disk began after electrode impedances fell below 25 K Ohms. Although the standard convention is to reduce impedance to 5 K Ohms or below (Teplan, 2002), the actiCHamp system uses active electrodes with noise reducing techniques built into the amplifier to ensure that impedances under 25 K ohms are sufficient for interpretable signals. Channels with impedance values above 25 K ohms were interpolated using data from neighboring electrodes with impedances below 25 K ohms. An auxiliary channel was used to record from a photosensor placed directly on a corner of the LED monitor. A 10-by-10 pixel square located under the photosensor was programmed to change from white to black during the onset of each visual stimulus; it changed from black to white during stimulus offset. Recording changes in screen luminance from the photosensor at 1 kHz allowed for precise timing of stimulus onset and offset with respect to the recorded EEG data.

### EEG Analysis

EEG signals were processed with BESA Research (v6.1) after re-referencing to a common average reference. First, notch (frequency 60 Hz, 2 Hz width) and bandpass (low cutoff 1 Hz, type forward, slope 6dB/oct; high cutoff 40 Hz, type zero phase, slope 12 dB/oct) filters were applied to all channels. Second, the signal on each channel was visually inspected to find, mark, and exclude the duration of all muscle artifacts. Third, a characteristic eye-blink was marked by finding an alternating deflection greater than 100 microvolts *μ*V) between the LOC and ROC signals. A template matching algorithm was then used to find all eye blink artifacts on all channels and remove the component of variance accounted for by the eye blinks (Picton et al., 2000; Ille et al., 2002). Finally, additional artifacts were isolated and excluded using amplitude (120 *μ*V), gradient (75 *μ*V), and low-signal (max. minus min) criteria (0.01). A participant’s data was used in further processing only if a minimum of 60% of trials survived this final artifact scan.

Following filtering and cleaning of EEG data, average evoked response potentials (ERPs) were computed for each condition (e.g., new, old (all), old_1_ (ST), old_2_ (LT), old_3_ (LT), and scrambled). The average ERPs for each condition were then used as input to group ERP statistical analyses performed with BESA Statistics v2.0 with appropriate multiple corrections across space and time (Maris and Oostenveld, 2007; Maris, 2012). Using this approach, statistical significance is assessed using nonparametric cluster permutation tests (N=1,000). Group ANOVAs were followed by pairwise t-test comparisons of different conditions in which contiguous clusters in space and time of coherent F (for ANOVA) or *t* values (for paired comparisons) exceeding an *a priori* corrected p-value of less than or equal to 0.05 were deemed significant. Summed F or t values of the clusters are compared to a null distribution of F or t sums of random clusters obtained by permuting the data order across subjects. This controls for type I errors due to multiple comparisons. The null hypothesis of the permutation test assumes that the assignment of the conditions per subject is random and exchangeable. The idea behind data clustering used in combination with permutation testing is that if a statistical effect is found over an extended time period in neighboring channels, it is unlikely that the effect occurred by chance. For t-values, a statistical effect can have a positive or negative direction and therefore positive and negative cluster values may be obtained. The positive or negative cluster value is the test statistic reported for each cluster, and the p-value reported is the one associated with that cluster based on permutation testing. For each of the 1000 permutations, new clusters are determined and the corresponding cluster values are derived for each cluster. Based on the new distribution, the alpha error of the initial cluster value can be directly determined. For example, if only 2% of all cluster values are larger than the initial cluster value, the initial cluster has a 2% chance that the null hypothesis was falsely rejected. This cluster would then be associated with a p-value of 0.02. The time with respect to stimulus onset and the sensor locations of each cluster are reported in addition to the cluster value and p-value (Table 1).

### Supervised Machine Learning

Supervised machine learning was performed on the cleaned and filtered ERP data matrix to investigate temporal and spatial patterns. Classification groups were new, old and scrambled scenes, and variances included between-subject variance. The first dimension of the input data matrix concatenated subject numbers for old, new, and scrambled condition. The second dimension was the number of channels used as the classification features. The third dimension was the number of time samples. The ERP matrix was segmented by a temporal sliding window using 50 ms length non-overlapping windows (Haufe et al., 2014). A classification pipeline was constructed which included a standardization scaler and linear stochastic gradient descent (SDG) classifier (Haufe et al., 2014; King and Dehaene, 2014). Cross validation was measured by a stratified 5-fold cross validation and data were shuffled within each fold. A total of 28 classifiers (each a 50 ms window) were cross-validated. Permutation tests were used to examine significance of the classifiers. The area under a receiver operating characteristic curve (ROC AUC) was used as the metric for the 2-class case of old and new. Accuracy was used as the metric for the 3-class case of old, new, and scrambled. Scalp topography patterns were computed as the dot product of the covariance matrix by the feature weights to visualize activity differences for new and old scenes (Haufe et al., 2014).

### MRI Scanning

Due to time constraints, the two subjects who underwent fMRI scanning completed only one 20- minute block of the VCRT. During the task, a T1-weighted anatomical MRI (TE 3.59 ms, TR 2000 ms, 9 degree flip angle, 249 mm field-of-view, 255 mm field-of-view, 1.00 mm slice thickness, 192 slices) and a single 20-minute EPI BOLD-fMRI run (800 timepoints, TE 30 ms, TR 1500 ms, 80 degree flip angle, 249 mm field-of-view, 3.10 mm slice thickness, 27 slices) were collected on a 3 tesla Siemens Prisma MRI scanner at the [Author University].

### fMRI Analysis

Alignment of the fMRI volumes, co-registration of each volume to the anatomical MRI, and statistical analysis of the fMRI timeseries was performed in AFNI Version AFNI_17.3.01 (Cox, 1996) using the *align_epi_anat.py* script and the *3dDeconvolve* command. The deconvolution of the responses to the jittered, randomized events (new, old, scrambled) assumed a hemodynamic response function, HRF(t), of int(g(t-s), s=0..min(t,d)), where g(t)=t^q*exp(- t)/(q^q*exp(-q)) and where t is time, d=1, and q=5. Statistical maps were computed using a general linear model with the six motion parameters as regressors of no interest and the new, old, and scrambled scene onset times as regressors of interest. General linear tests included old vs. new, old vs. scrambled, and new vs. scrambled. Pial layer and inflated cortical surfaces were made using Freesurfer v5.3 (Fischl, 2012) and used to display single-subject statistical maps in AFNI’s SUMA viewer.

## Results

A list of all statistical comparisons and p values obtained from the behavioral, EEG (including standard ERP and supervised machine learning), and fMRI analyses is included in Table 1.

### Behavioral

Subjects performed well above chance (50%) discriminating old from new stimuli (85.7% correct, S.D. 8.5, Fig 2a). When old scene recognition was analyzed as a function of the three time intervals, accuracy was best for the short-term (ST) interval and declined at each of the two longer-term (LT) intervals (repeated measures ANOVA, F_(2,56)_=186.3, p<0.001, Fig 2a). A similar pattern was obtained when the dependent measure was sensitivity (d-prime) instead of percent correct (repeated measures ANOVA, F_(2,54)_=165.3, p<0.001, Fig 2b). Subjects responded faster to old scenes compared to new scenes (old=967.6 ms, s.d. 58.95 vs. new=1012.1 ms, s.d. 79.45, paired t_(28)_=4.067, p<0.001, Fig 2c). A signal detection breakdown of response times confirmed differences among hits (957.5 ms, s.d. 57.62), correct rejections (986.9 ms, s.d.81.80), misses (1040.5 ms, 92.59 s.d.) and false alarms (1352.1 ms, 173.73 s.d.). A repeated measures ANOVA to compare the effect of type of response on the dependent variable response time was significant (F_(3,84)_=86.63, p<0.001, Fig 2c). Post-hoc tests revealed that hits were significantly faster compared to both false alarms (post-hoc t=-10.946, p_bonf_<0.001) and misses (post-hoc t=-5.845, p_bonf_<0.001) but not correct rejections (post-hoc t=-2.419, p_bonf_=0.134).

**Figure 2.**
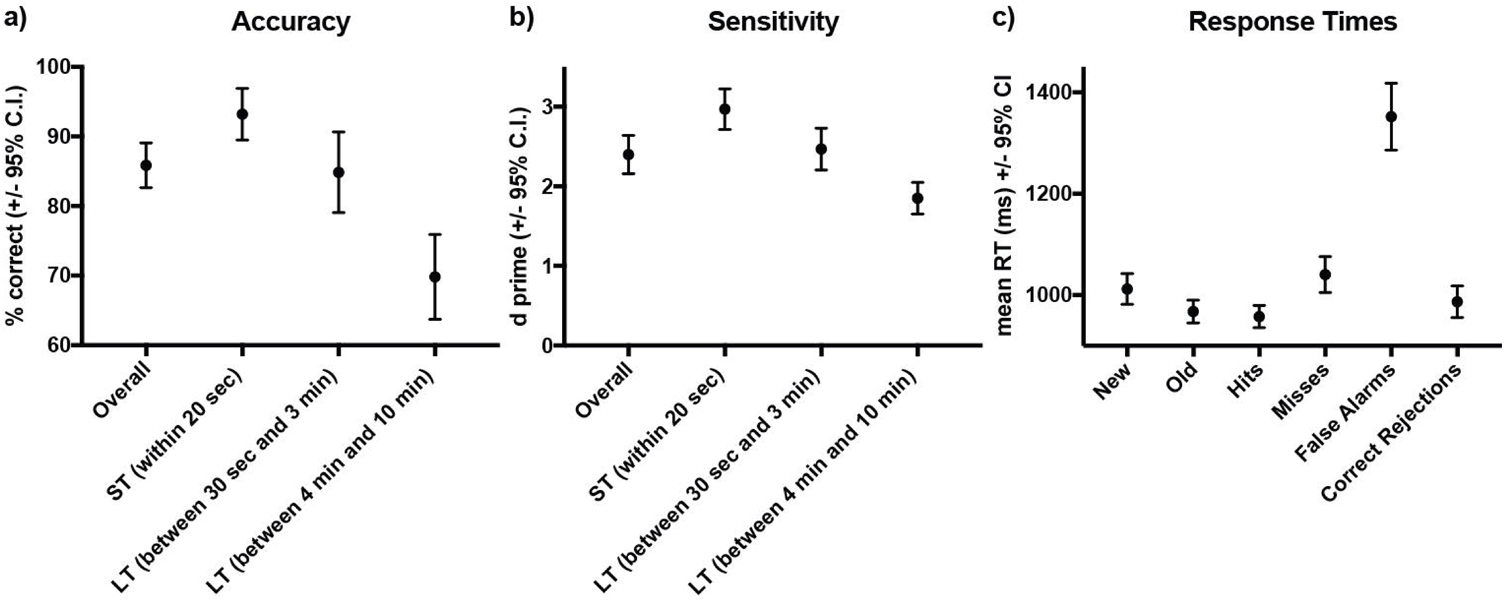
Behavioral Accuracy and Response Times during Scene Recognition. Overall percent correct for new/old recognition was above chance (50%), was highest for the short-term interval and declined at the two longer term intervals (a). A similar pattern was found for sensitivity (b). Response times were longer for new compared to old scenes, fastest for hits and correct rejections, slower for misses, and slowest for false alarms (c).

### Evoked Response Potentials

#### ANOVA New vs. Old vs. Scrambled ERPs

A group ANOVA of the ERP data with three levels (new, old, and scrambled conditions) produced a highly significant, extended cluster (cluster value=641760, p<0.001) ranging from 59 ms to 1399 ms post-stimulus. This indicated that evoked responses to the three stimulus types were distinguishable through most of the post-stimulus time window. The scalp topography pattern at early and later post-stimulus times indicate higher magnitude positivities in parietal sensors (Fig 3a,b) and negativities (Fig 3c,d) in frontal sensors for old scenes.

**Figure 3.**
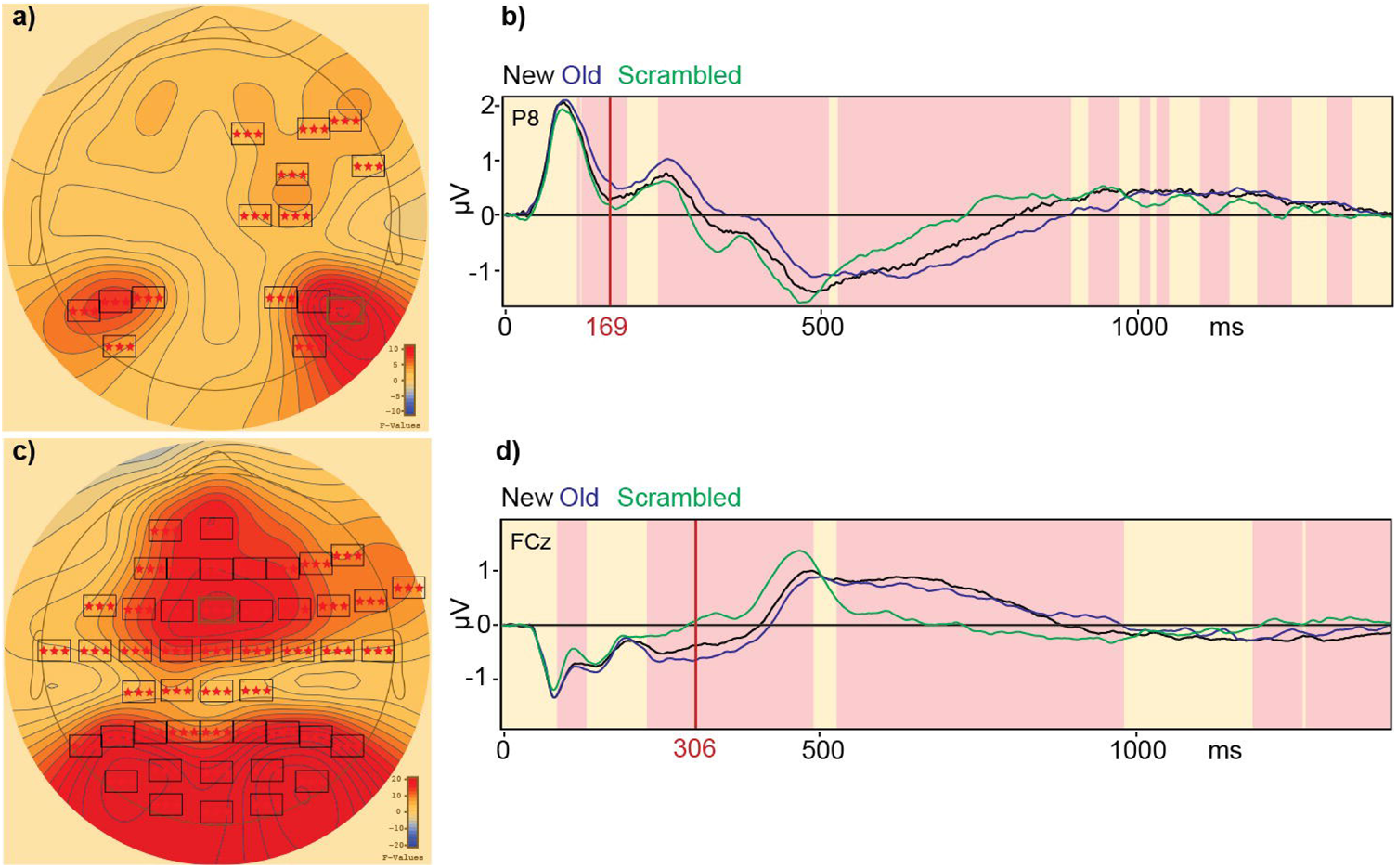
Evoked Response Potentials Distinguish New, Old, and Scrambled Scenes. A group ANOVA of ERPs distinguish the three scene types - new, old, and scrambled – across most of the stimulus window. The early scalp topography, especially at parietal sensor P8, showed higher magnitude positivities (a,b). Later in time a more negative deflection in a centro-frontal sensor FCz characterized the response evoked by new and old scenes (c,d) compared to scrambled scenes (c,d). Boxes with colored asterisks in a and c indicated sensors that are part of significant spatiotemporal clusters, while red shaded areas in b and d indicate time intervals of significant differences in group ERPs (p<0.01).

After this ANOVA, post-hoc paired comparison t-tests were computed to compare the evoked response to old versus scrambled conditions and also to compare the responses to new versus the scrambled conditions. Early responses (∼200 ms) in parietal sensors showed greater positivities for old scenes compared to scrambled scenes (Fig 4a,b). Later responses (∼300 ms) in centro-frontal sensors showed more negative evoked responses to new scenes compared to scrambled scenes (Fig 4c,d).

**Figure 4.**
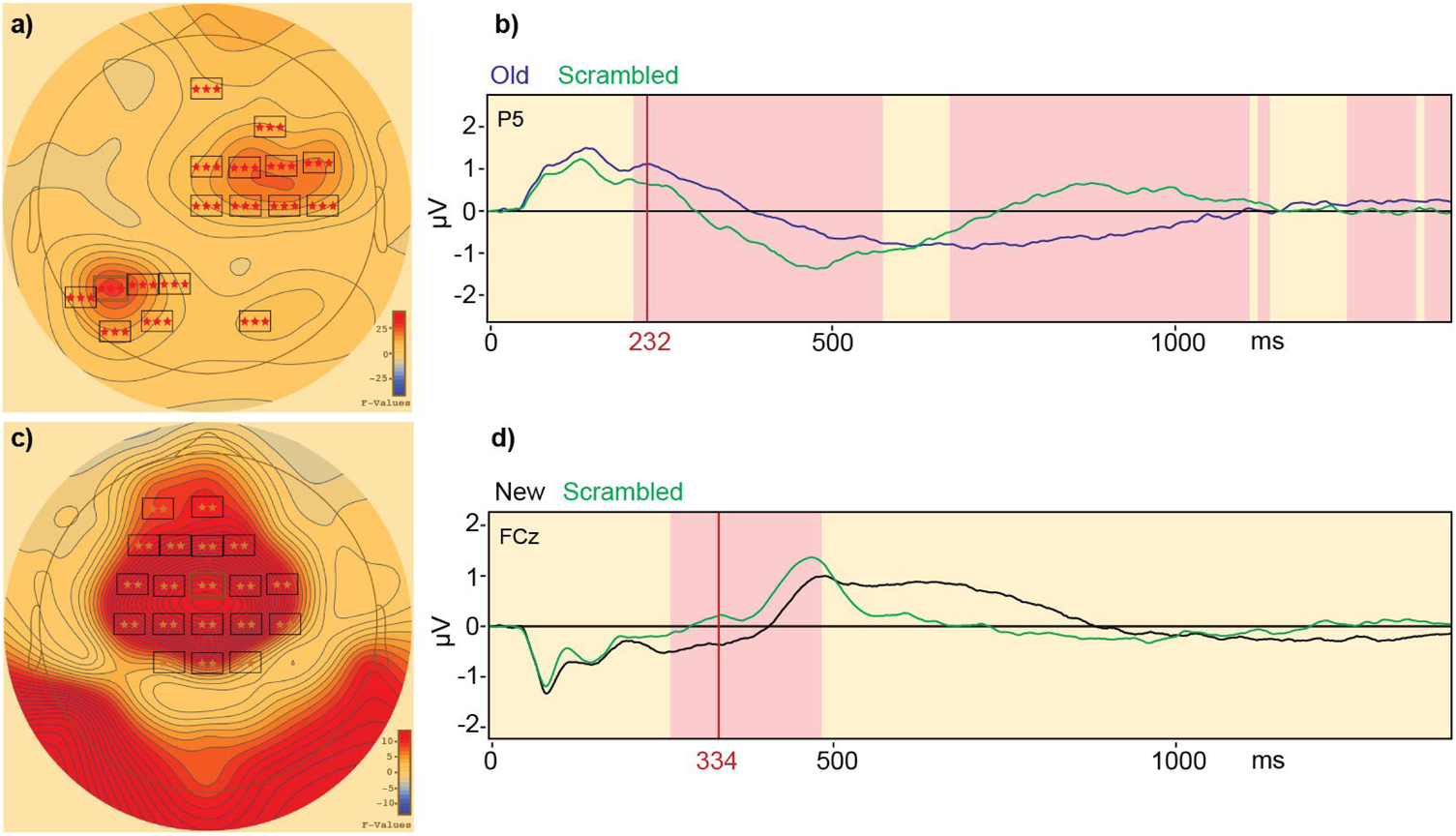
Evoked Response Potentials Distinguish Old from Scrambled and New from Scrambled Scenes. Post-hoc paired comparison t-tests computed after the three level ANOVA compared evoked responses show that early responses (∼200 ms) in parietal sensor P5 were characterized by greater positivities for old scenes compared to scrambled scenes (a,b). Later responses (∼300 ms) in the centro-frontal sensor FCz showed more negative evoked responses to new scenes compared to scrambled scenes (c,d). Boxes with colored asterisks in a and c indicated sensors that are part of significant spatiotemporal clusters, while red shaded areas in b and d indicate time intervals of significant differences in group ERPs (p<0.01).

#### Paired T-Test of New vs. Old ERPs

To determine whether evoked responses differed for new and old scenes, a paired t-test of the group ERP data was computed and showed that parietal ERPs were positive and greater for old versus new scenes by around 200 ms (cluster value=-25074.4, p<0.001, 189 ms to 750 ms, Fig 5a,b). It was also found that evoked responses in a fronto-temporal cluster were positive and greater for new versus old scenes (cluster value=16478.3, p<0.001, 194 to 744 ms, Fig 5c,d).

**Figure 5.**
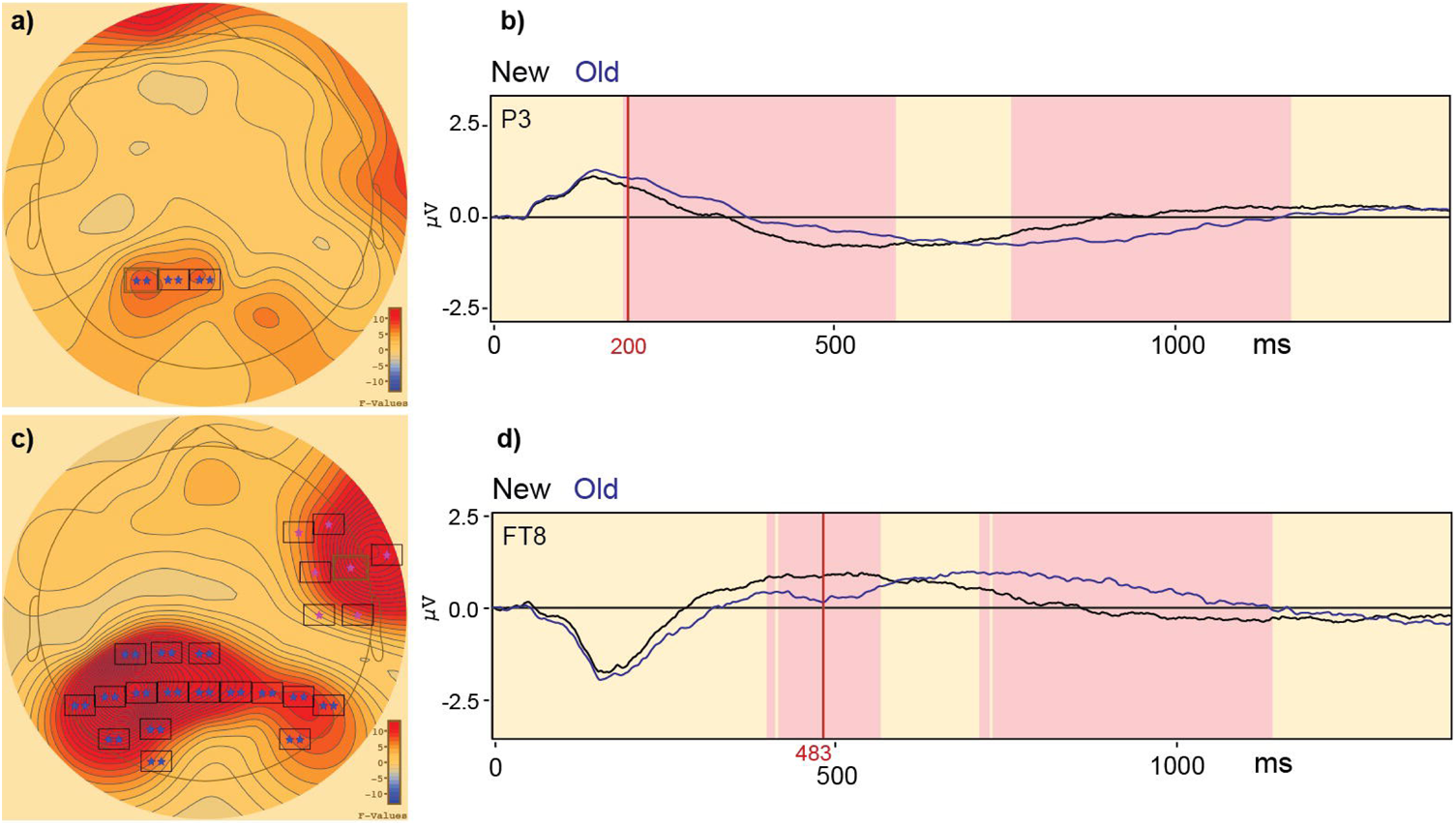
Differences in Evoked Response Potentials for New and Old Scenes. A paired t-test of the group ERP data revealed parietal sensors with greater positivities for old compared to new scenes (a,b). A fronto-temporal cluster showed greater positivities for new compared to old scenes. Boxes with colored asterisks in a and c indicated sensors that are part of significant spatiotemporal clusters, while red shaded areas in b and d indicate time intervals of significant differences in group ERPs (p<0.01).

#### ANOVA of Short- versus Long-Term Old Scene ERPs

Motivated by the behavioral results showing different levels of recognition performance depending on whether a scene was first presented in a short- or long-term time interval, a group ANOVA of the sensor level ERPs was computed to determine whether and where evoked responses could distinguish the three types of old scenes, short-term (old_1_ within 20 sec), intermediate (old_2_ between 30 sec and 3 minutes), and long-term (old_3_ between 4 and 10 minutes). Four clusters were found indicating discrimination as a function of retention interval. In descending order of magnitude, these included a cluster with greater positivities evoked by old scenes in the longer-term intervals compared to the short-term interval (cluster value=9621.85, p=0.001, 355 to 517 ms, peak at 454 ms at sensor FC1, old_1_ 0.1917 vs. old_2_ 0.4522 vs. old_3_ 0.5099, Fig 6c,d). The second cluster consisted of more negative responses evoked by old scenes in the longer-term intervals compared to the short-term interval (cluster value=9128.21, p=0.001, 364 to 554 ms, peak at 456 ms at sensor P8, old_1_ −0.4623 vs. old_2_ −0.6836 vs. old_3_ − 0.7497). The third cluster consisted of greater positivities evoked by old scenes in the longer-term intervals compared to the short-term interval (cluster value=4082.77, p=0.004, 228 to 324 ms, peak at 291 ms at sensor TP8, old_1_ −0.1368 vs. old_2_ 0.1751 vs. old_3_ 0.0775, Fig 6a,b). The fourth cluster consisted of a greater positivity evoked by old scenes in the short-term interval compared to the longer-term intervals (cluster value=3437.09, p=0.012, 259 to 326 ms, peak at 287 ms at sensor C3, old_1_ 0.1109 vs. old_2_ −0.1929 vs. old_3_ −0.0141).

**Figure 6.**
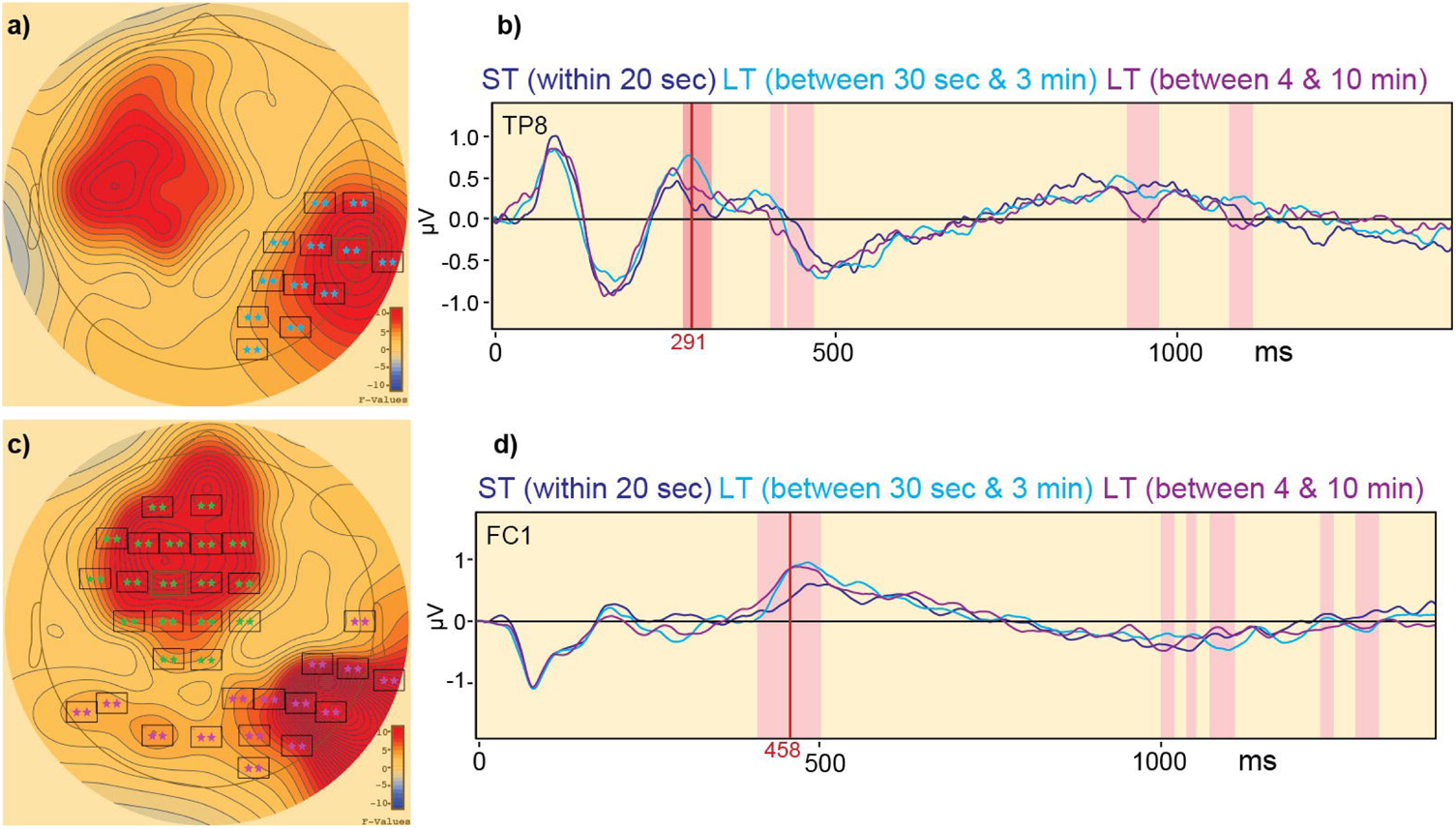
Frontal and Parietal ERPs Distinguish Old Scenes Presented at Long- versus Short-Term Intervals. A difference in ERPs centered at electrode TP8 occurring 291 ms after stimulus onset (a) shows the greatest positivity for old scenes first presented within 30 sec to 3 minutes (b, light blue) and the least positivity for old scenes first presented with 20 sec ago (b, purple). A difference in ERPs centered at electrode FC1 occurring 458 ms after stimulus onset (c) shows greater positivities for old scenes first presented between 30 sec and 10 minutes ago (d, blue and light purple) and the least positivity for old scenes first presented 20 sec ago (d, purple). Boxes with colored asterisks in a and c indicated sensors that are part of significant spatiotemporal clusters, while red shaded areas in b and d indicate time intervals of significant differences in group ERPs (p<0.01).

### Supervised Machine Learning

A supervised machine learning algorithm was used to determine how accurately scene type could be decoded as a function of time after stimulus presentation. Decoding of stimulus type in a 2-class scenario with new and old scenes (Fig 7a) as well as in a 3-class scenario with new, old, and scrambled scenes (Fig 7b) reached levels significantly above chance by the 275 ms post-stimulus time interval. Decoding accuracy peaked in a time interval from 900 to 1100 ms post-stimulus, overlapping with the range of average subject response times (Fig 2c). When the decoding results were plotted as a function of post-stimulus time, early posterior activity for new scenes was visualized from 50 to 200 ms and then there was a switch to old scenes from 200 to 600 ms (Fig 8).

**Figure 7.**
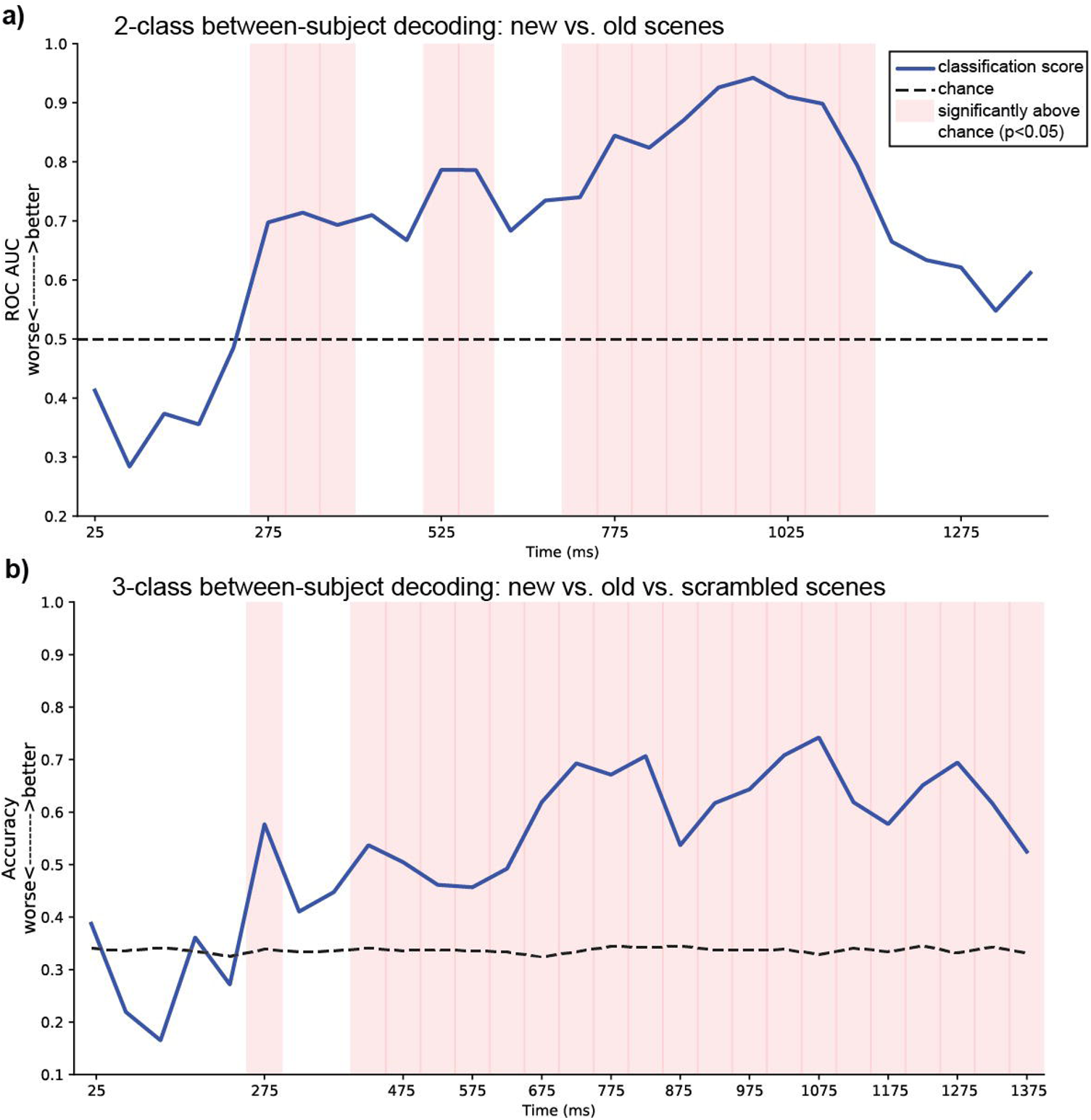
Decoding of Scene Type with Supervised Machine Learning. Classification of stimulus type using supervised machine learning in a 2-class scenario with new and old scenes (a) and in a 3-class scenario with new, old, and scrambled scenes (b) reached levels above chance (red shaded bars) by the 275 ms post-stimulus time interval.

**Figure 8.**
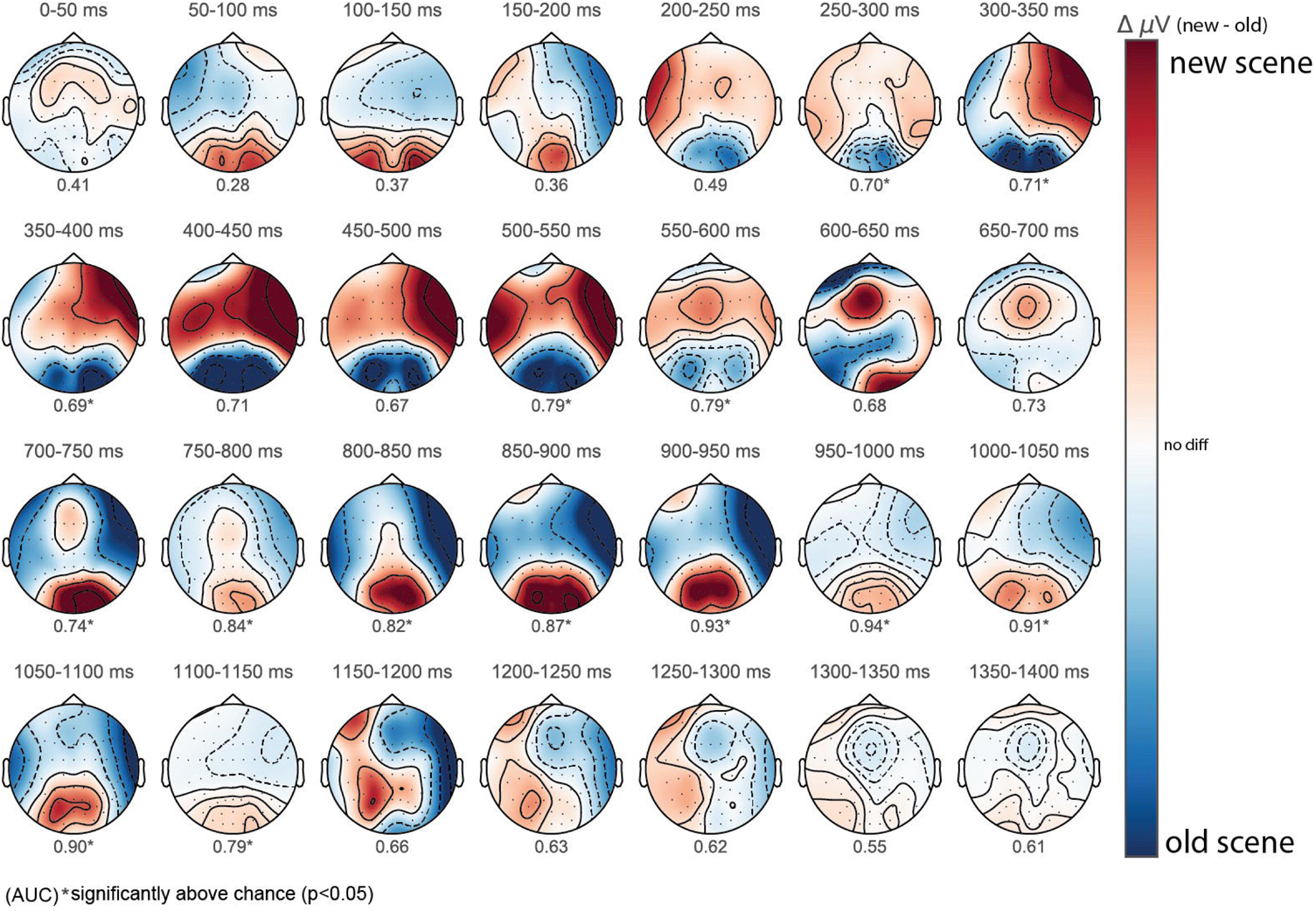
Spatiotemporal Scalp Topography Patterns from Machine Learning Classification Show Widespread Contributions to Decoding Scene Type. The decoding results were plotted as a function of post-stimulus time interval to visualize differences for new and old scenes. Early posterior activity for new scenes dominates from 50 to 200 ms and then there is a switch to old scenes from 200 to 600 ms.

### Functional MRI

Single-subject fMRI was used to examine activity differences evoked by presentation of the old and new scenes in two subjects. T-maps (alpha threshold at p<0.01, false discovery rate q<0.05, cluster extent threshold of 100 voxels) reflecting a contrast comparing old versus new scenes produced four separate clusters in each of the two subjects. In the first subject (Fig 9a,b) all four clusters were characterized by greater activity for old compared to new scenes. The four clusters in descending order of size were 1) right precuneus (999 voxels with peak at +5, −45, +40 mm MNI space), 2) right superior parietal lobule/angular gyrus (959 voxels with peak at +39, −66, +49 mm), right middle frontal gyrus/BA6 (130 voxels with peak at +38, +22, +43), and 4) right middle frontal gyrus/BA10 (118 voxels with peak +30, +68, +2). Four clusters were also found in the second subject (Fig 9c,d), and were also characterized by greater activity for old compared to new scenes. In descending order of size they included 1) left superior frontal gyrus (943 voxels with peak at −24, +2, +77 mm), 2) right superior parietal lobule/angular gyrus (193 voxels with peak at +40, −48, +65 mm), 3) right inferior frontal gyrus pars opercularis BA45/47 (119 voxels with peak at +57, +19, −4 mm), and 4) right superior medial frontal gyrus (117 voxels with peak at +4, +55, +49 mm).

**Figure 9.**
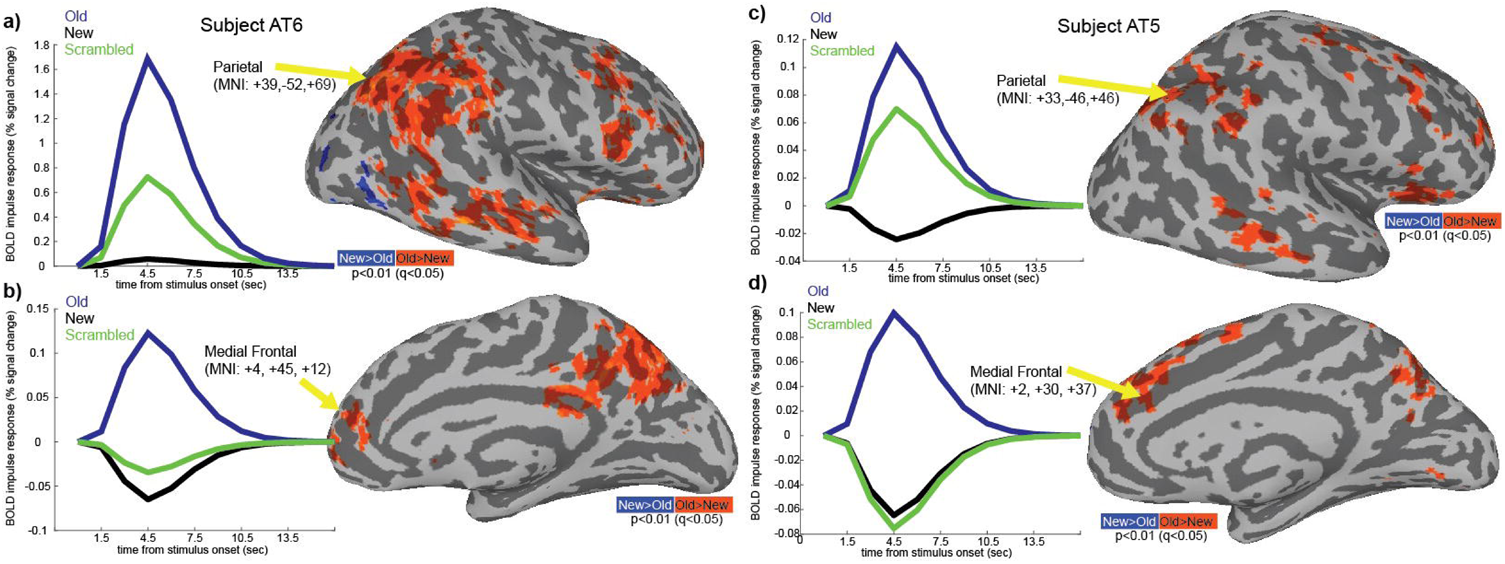
Single-Subject fMRI Reveals Widespread Cortical Contributions to Scene Memory. In two subjects, a contrast of old and new scenes produced activity differences dominated by more activity for old compared to new scenes. In subject 1 (a,b) all four clusters were found in right hemisphere and included precuneus, angular gyrus, and middle frontal gyrus (BA6 and BA10). In subject 2 (c,d) three out of four clusters were found in the right hemisphere included left superior frontal gyrus, superior parietal lobule (BA 40/7), and superior medial frontal gyrus (BA8). The peak of the cluster of activity in left hemisphere of subject 2 (not shown) was located in superior frontal gyrus (BA 6).

## Discussion

The remarkable human capacity for detailed scene recognition memory has been extensively documented in previous behavioral studies, yet the neural bases supporting this ability remain to be fully understood. Previous neural studies have focused mostly on understanding the basis for scene specificity and therefore have utilized designs in which categorization is the required cognitive task for making decisions about stimulus sets consisting of scenes and other complex visual stimuli like faces, animals, or objects or scenes with faces, animals or objects (Thorpe et al., 1996; Tsivilis et al., 2001; Rousselet et al., 2002; Rousselet et al., 2004; Harel et al., 2016). Understanding categorization ability, although certainly an interesting and highly-developed cognitive function, was not the focus of the present study. Instead, the questions addressed here involved scene memory. The primary object was to understand when and how neural patterns distinguish novel, familiar, and scrambled scenes. We therefore included as a baseline condition a set of phase-scrambled scenes in which color and spatial frequency were similar to the real outdoor color scenes. Subjects could not, however, infer from the phase-scrambled scenes anything about place, spatial layout, or meaning from the content of the images. The use of scenes and phase-scrambled counterparts, rather than complex stimuli of different categories, makes the proactive interference experienced during viewing of the interspersed scrambled scenes perceptual rather than categorical.

The first novel contribution of the present study is the characterization of the spatiotemporal neural patterns associated with distinguishing new and old scenes from the phase-scrambled versions. A group ANOVA revealed parietal and frontal ERPs discriminated the three scene conditions (i.e., new, old, and scrambled) as early as 59 ms after stimulus onset. Inspection of the early evoked patterns in Fig 3d shows that this involved a greater negativity for new and old scenes compared to the scrambled set in a centro-frontal region. When post-hoc direct comparisons of different scene types were made to the scrambled set, parietal positivities were greater for old scenes by 210 ms (Fig 4a,b) and centro-frontal negativities were greater for new scenes by 235 ms (Fig 4c,d). The greater parietal positivity at 210 ms is consistent with recent finding that a P2 amplitude peaking at 220 ms is sensitive to distinguishing open and closed natural scenes (Harel et al., 2016). Intuitively the pattern of neural results with fastest evoked responses for scrambled scenes followed by longer times for real scenes makes sense since identification of a stimulus as a scrambled scene (Fig 1a) happens quickly at a perceptual level; identifying aspects of a non-scrambled scene that include the content conveying meaning, place and layout relating to the real world likely involves additional neural computations.

The second novel contribution of the present study is the characterization of the temporal dynamics associated with discriminating old and new scenes. Direct paired comparisons of evoked responses showed elevated parietal ERPs for old compared to new scenes by 189 ms (Fig 5a,b). For new scenes, fronto-temporal ERPs were less negative by 194 ms and eventually developed a greater positivity relative to old scenes. The pattern of findings obtained is reminiscent of previously reported old/new effects found with other recognition paradigms (Sanquist et al., 1980; Warren, 1980; Wagner et al., 2005; Rugg and Curran, 2007). The old/new effect has been described as the more positive-going evoked response to old (i.e., studied) items compared to new (i.e., unstudied items). The effect is usually seen as a left-sided parietal response peaking 400 to 500 ms after stimulus onset. In the present study, we employed a rapid jittered design with stimulus presentation rate of about 6.6 stimuli every 20 seconds and, because of the speeded presentation, did not attempt to have subjects rate familiarity strength using a remember/know procedure after each scene presentation. This means that we cannot determine whether the parietal responses evoked by old scenes were enhanced for scenes actually remembered compared to scenes merely recognized as familiar (Warren, 1980).

The third novel contribution of the present study is the characterization of differences in evoked responses to old scenes as a function of retention interval. The study was designed so that some scene presentations were repeated a second time within a 20 sec window after the first presentation. We labeled this second presentation as occurring within a short-term interval since 20 sec is often assumed as the temporal limit for short-term memory based on classic interference paradigms (Peterson and Peterson, 1959; Keppel and Underwood, 1962). Although it should be noted that while this assumption is based on paradigms that assess memory based on verbalizable items like letter trigrams, it has been recently demonstrated using time-frequency that the right parietal region is active during the maintenance (6 sec delay) of two scenes in short-term memory (Ellmore et al., 2017). Two other intervals of between 30 sec and 3 min, and between 4 and 10 min were classified as long-term intervals. The behavioral results support a distinction among these three intervals with accuracy highest for short-term recognition and falling significantly for the later intervals, but remaining well-above chance (Fig 1a,b). The evoked neural patterns obtained in the present study also support a distinction between short- and long-term scene memory. For old scenes presented after a long-term interval, parieto-temporal and centro-frontal ERPs were greater by 228 and 355 ms, respectively, compared to old scenes presented after a short-term interval (Fig 6). This finding is consistent with recent work showing a rapid and independent role for parietal cortex in a wider network for developing longer-term memory (Brodt et al., 2016).

The main conclusions of this study are based on analyses of the primary data, which include EEG data collected from 27 subjects. We conducted secondary analyses and collected other hemodynamic data that bolster some of the conclusions about the evoked spatiotemporal patterns. First, we performed supervised machine learning to build a model that makes predictions based on evidence in the presence of uncertainty. Using this model, we could identify patterns across the feature set of all EEG sensors to make predictions in time about which stimulus class (i.e., new, old, or scrambled) the subjects have been presented. The machine learning results demonstrated an ability to distinguish among the three scene types by 275 ms (Fig 7) after stimulus onset. Changing patterns of activity particularly in parietal-occipital and fronto-temporal regions evoked by the old and new scenes were similar for machine learning (Fig 8) compared to the scalp topography maps generated by conventional ERP analysis (Fig 3 to 6). We can conclude from these results that the changing pattern of distributed activity across the scalp can be used to decode old from new scenes, and also classify old and new from scrambled scenes.

Second, while the EEG reflects dipoles from neural activity in the brain measured by sensors located on the scalp, there is considerable uncertainty about where these signals originate. Single-subject fMRI however allows for more precise spatial localization by obtaining a hemodynamic measure of the elevated blood oxygenation levels occurring as a result of increased local field potentials. While temporal evolution of the BOLD signal is lagged by as much as 6 sec from stimulus onset, and therefore the temporal resolution is far inferior to that obtained with EEG, we collected fMRI data in two subjects who performed the same task but completed only one fourth of the number of trials as compared to the subjects who completed the EEG experiments. The fMRI results confirm the general pattern at the EEG sensor level that scene memory involves widespread cortical areas including medial frontal, parietal, and temporal regions. We found a consistent pattern of fMRI activity across the subjects that was greater overall for old compared to new scenes (Fig 8) in parietal, temporal, and medial frontal regions. Unfortunately, the limited temporal resolution of fMRI does not allow us to distinguish differences in activity during narrow post-stimulus time windows of between for example 200 and 600 ms, which EEG can resolve quite easily using jittered rapid serial visual presentation. Because of the fewer trials obtained in the subjects who completed fMRI, we did not attempt to distinguish differences for old scenes presented after long-compared to short-term intervals. An additional limitation in the fMRI data is that, in order to maintain a fast repetition time, we traded off total slice numbers in the axial acquisitions. Thus, we did not capture fully the basal and medial temporal areas including hippocampus and parahippocampal gyrus. Therefore, in this limited sample, we cannot investigate neural patterns in these regions during scene recognition.

In conclusion, we present converging evidence from multiple modalities and analysis approaches that the high capacity human scene recognition memory system is supported by neural activity patterns occurring by 200 ms in widespread frontal, temporal and parietal cortical regions. Changes occurring later, between 225 and 550 ms, allow a distinction between scenes first presented 20 sec ago compared to several minutes ago. These findings provide a baseline by which to evaluate in future neural studies the more nuanced aspects of the scene memory system, including how scene information is consolidated rapidly and available for accurate recognition after even longer retention intervals, including days and beyond (Chandler, 1991), and how neural patterns resist accumulating proactive interference (Makovski and Jiang, 2008) as hundreds or even thousands more scenes are encoded for subsequent recognition.

## Table Legends

**Table 1. Statistical Table.** The statistics and associated p-values are reported for each of the behavioral, group ERP, machine learning, and fMRI analyses. Sensor locations are named according to the extended international 10/20 system. Means for ERP conditions are in units of microvolts and are integrated across the entire temporal window indicated by start and end time. For machine learning accuracy is reported for 3-class decoding, while area under the curve is reported for 2-class decoding. Cluster sizes for fMRI are in number of voxels where each voxel represents 3.134×3.134×3.100 mm^3^. SPL=superior parietal lobule, MFG=middle frontal gyrus, IFG=inferior frontal gyrus, Sup. MFG=superior medial frontal gyrus.

## Author Contributions

TME: conceptualization, funding acquisition, project administration, resources, supervision, visualization, formal analysis, writing (original draft), writing (review and editing).

CPR, KN, and NM: data curation, investigation, formal analysis, visualization, writing (original draft), writing (review and editing).

## Acknowledgements

The authors would like to thank Dr. A. Duke Shereen of the CUNY ASRC MRI Facility for help with fMRI data acquisition.

